# Finding missing links in interaction networks

**DOI:** 10.1101/695726

**Authors:** J. Christopher D. Terry, Owen T. Lewis

**Affiliations:** Department of Zoology, University of Oxford, Oxford, OX1 3PS, UK

**Keywords:** ecological network, bipartite network, missing links, sampling completeness, network metrics

## Abstract

Documenting which species interact within ecological communities is challenging and labour-intensive. As a result, many interactions remain unrecorded, potentially distorting our understanding of network structure and dynamics. We test the utility of four structural models and a new coverage-deficit model for predicting missing links in both simulated and empirical bipartite networks. We find they can perform well, but that the predictive power of structural models varies with the underlying network structure. Predictions can be improved by ensembling multiple models. Sample-coverage estimators of the number of missed interactions are highly correlated with the number of missed interactions, but strongly biased towards underestimating the true number of missing links. Augmenting observed networks with most-likely missing links improves estimates of qualitative network metrics. Tools to identify likely missing links can be simple to implement, allowing the prioritisation of research effort and more robust assessment of network properties.

## Introduction

Networks documenting trophic, competitive or mutualistic interactions among species are an essential tool for understanding the structure and dynamics of ecological communities (Memmott 2009, Poisot et al. 2016b, Dormann et al. 2017, Delmas et al. 2019). Interaction networks can be a better indicator of the resilience of communities than species counts (Valiente-Banuet et al. 2015) and are increasingly applied in conservation (Tylianakis et al. 2010) and biomonitoring (Gray et al. 2014).

Documenting the species present in a system can be difficult enough (Gotelli and Colwell 2010); accurately enumerating their interactions is even more challenging. As a result, ecological interaction networks are routinely under-sampled, with consequences for our understanding of communities (Goldwasser and Roughgarden 1997, Blüthgen 2010, Chacoff et al. 2012, Jordano 2016). Ecological networks are sparse and hence any unobserved interactions could be either ‘missing links’ (interactions that would be observed with complete sampling), or true negatives. Differences or deficiencies in sampling can lead to divergent conclusions about underlying ecological mechanisms (Lee and Guénard 2019). Furthermore, under-sampling will skew the apparent prevalence of interactions towards frequent interactions and may miss infrequent interactions entirely (Dormann et al. 2017).

Common solutions to this problem have included rarefying networks to a consistent level of sampling (Morris et al. 2014), sampling until metric values stabilise (Rivera-Hutinel et al. 2012, Costa et al. 2016), explicit observation models (Weinstein and Graham 2017), and the use of network metrics that are comparatively robust to under sampling (Vizentin-Bugoni et al. 2016). There has been an extensive literature testing these different approaches (Nielsen and Bascompte 2007, Blüthgen et al. 2008, Poisot et al. 2012b, Fründ et al. 2016, Henriksen et al. 2019, de Aguiar et al. 2019), often arguing for an emphasis on quantitative rather than qualitative metrics, which reduce the influence of missed infrequent interactions.

Nonetheless, infrequent interactions may play an important role in ecological communities. Since networks are not static, there is value in distinguishing infrequent interactions from those that are genuinely absent. Rarely-observed links have the potential to grow in importance with changes in biotic (Terry et al. 2017) or abiotic conditions (Staniczenko et al. 2017). At equilibrium, interactions may occur at low frequency, yet be dynamically important. For instance, while interactions involving a species suppressed to a low level by a consumer may be observed infrequently, removal of the consumer can allow prey abundance to increase rapidly (Paine 1980). At a network level, the prevalence and distribution of apparently weak or infrequent interactions can have marked consequences for network stability and dynamics (McCann et al. 1998, Berlow 1999, Gellner and McCann 2016, Jacquet et al. 2016, Kadoya et al. 2018).

Ecological networks are manifestly non-random (Dunne and Pascual 2006, Montoya et al. 2006, Vázquez et al. 2009, Bascompte and Jordano 2013). In principle, it should be possible to use this underlying structure to help infer where links are most likely to have been missed (Bartomeus 2013, Morales-Castilla et al. 2015, Valdovinos 2019). To this end, a number of missing link inference approaches have been tested on ecological networks. These have included hierarchical structuring models (Clauset et al. 2008), linear filtering (Stock et al. 2017), matching-centrality models (Rohr et al. 2016), stochastic block models (Guimera and Sales-Pardo 2009) and *k*-nearest neighbour recommenders (Desjardins-Proulx et al. 2017). These models have shown that missing links can be inferred from partially-observed structured networks. However, although ecological networks are used as part of benchmarking tests, often the inference scenarios are not orientated towards ecological needs and to date there has been limited uptake of these methods from the ecological community.

Furthermore, it is common in empirical ecological networks for species to have been sampled to markedly different extents (although see Novotny *et al.* 2010). This has the potential to give further information about the potential location of missing links, since missing links are more likely for poorly-sampled species (Blüthgen et al. 2008).

Here, we explore the potential to identify missing links in a wide range of ecological bipartite networks with a range of models, including a new species-level coverage-deficit approach which incorporates information on the completeness of sampling for individual species. We show that inferential models can perform well, especially when used in ensemble, and demonstrate how they could inform empirical work to document ecological networks and help evaluate hypotheses.

## Methods

First we present a set of predictive models to detect missing links and introduce a suite of simulated and empirical networks. These are used to explore each model’s capacity to identify missing links and improve the estimation of network-level metrics.

### Predictive Models

Our predictive models seek to identify the true missing links within an observed bipartite adjacency matrix **O** composed of observation counts *o_ij_*. This has rows representing each focal layer species *i* and columns detailing each potential interaction partner *j*. We assume that this set of observations is derived from a set of true interaction frequencies, where the relative rate of interaction between two species is given by *a_ij_*. We use our statistical models to assign to each unobserved interaction in a network a relative probability of being a ‘missing link’ (false negative), rather than a true negative: *P*(*a_ij_* > 0 | **O**) for the cases where **O_ij_** = 0. We assume throughout that the species observed (the dimensions of **O**) are a complete list – we do not attempt to predict the identity of interactions with unobserved species. Here we introduce and detail a new ‘coverage deficit’ model and four structural models described in the literature. All models were fit using the *cassandRa* R package (Terry 2019). Example predictions on a simulated network of each of the core models are shown in Figure 1.

**Figure 1.**
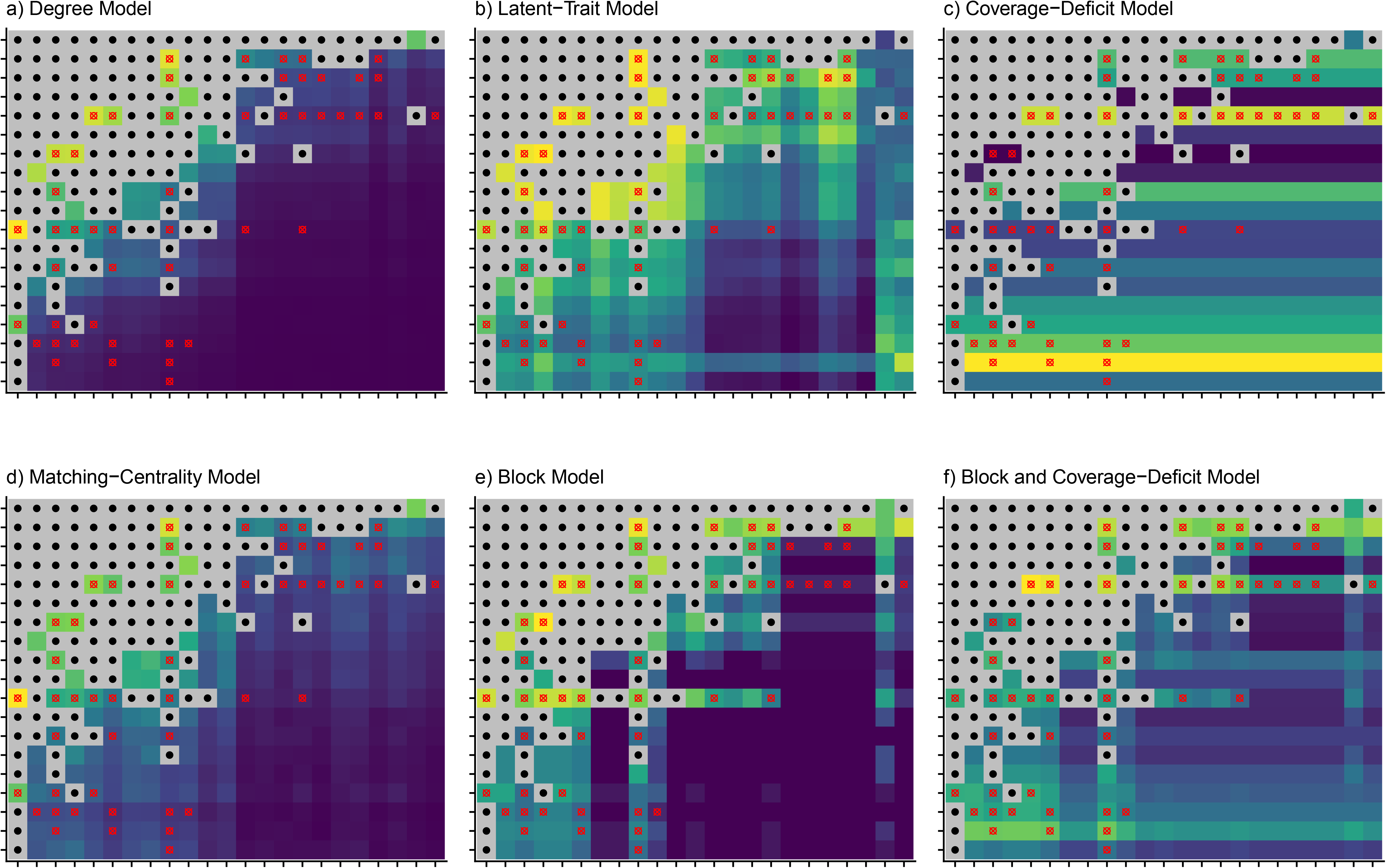
Demonstration of the fits of predictive models to a subsampled simulated bipartite network. Brighter colours indicate a higher probability of an unobserved interaction being present according to that model. Black dots (●) indicate observed interactions and red targets (□) indicate unobserved missing links that we wish to identify. AUC values for this example were as follows: Degree: 0.80, Latent-Trait: 0.76, Matching-Centrality: 0.81, Block: 0.88, Coverage-Deficit: 0.7, Averaging the Block and the Coverage-Deficit model 0.92.

### Sample size and Coverage-Deficit Models

Interactions are more likely to have been missed between species that are relatively poorly sampled. When ecological bipartite networks are constructed, it is common that one level (the focal level) of a network is sampled directly and the interaction partners in the other level counted. Examples include gathering insect hosts and rearing parasitoids from them, or observing pollinators that visit a set of plant species. This sampling may be opportunistic, matched to the local abundance, or standardised in some other manner. The result is a set of observed interaction counts with each focal layer species. Although interaction counts may be later rescaled to better represent local abundances, here we assume the raw observation data are available of discretisable relationships where individual interaction events can be observed. This includes many ecological interactions such as parasitism, pollination visits and feeding observations, but excludes diffuse facilitatory or competitive interactions. A simple approach assigns a relative probability of missed interactions based solely on the number of observations of each focal layer-species:

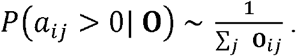

We refer to this as our ‘sample-size’ model.

This approach can be refined using techniques from the extensive literature developed to assess the sample completeness of species inventories using information about the most infrequent observations to estimate the number of missing observations (Chao and Jost 2012). Rooted in information theory, these approaches can be applied to any case of estimating sample coverage, including interaction sample coverage (Chacoff et al. 2012, Jordano 2016). The core of our coverage-deficit model estimates the interaction partner sample completeness for each focal layer species, based on the observed occurrence frequencies and the ‘Chao1’ estimator (Chao 1984, Chao & Jost, 2012). Where data is sufficient, this coverage deficit estimator (*Ĉ_def_*) is an estimate of the proportion of interactions that occur between as-yet-unobserved partners, or alternatively the probability that the next interaction partner observed would not yet have been observed interacting with that species. It is defined as:

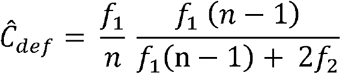

where *f*_1_ is the number of interactions observed only once, *f*_2_ is the number of interactions observed exactly twice, and *n* is the total observed interactions involving that focal species. For each focal layer species *i*, we use a *Ĉ_def_i__* estimate to assign a relative probability to each potential missing interaction as:

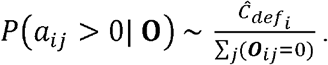

Unfortunately, ecological interaction networks typically contain a number of focal layer species that are extremely poorly observed, resulting in situations where the Chao1 estimator for coverage deficit is undefined or inappropriate. For very poorly sampled species, the distribution of interactions cannot itself provide much information (Colwell and Coddington 1994). Where observations are exclusively singletons (*f*_1_ = *n*) the index will estimate zero coverage. At the other end of the scale, if a focal species is observed with no singleton interactions the index will estimate that there is zero sample coverage deficit.

In these cases, we use a simple binomial model to estimate *Ĉ_def_*, the probability that the next interactor drawn will not yet have been observed interacting with the focal species. The likelihood of not having yet observed any of the unobserved missing links is: *L* = (1 − *Ĉ_def_*)^*n*^. While the maximum likelihood estimate of *Ĉ_def_* will be 0, with a Bayesian approach and assuming a flat prior the posterior mean can be found directly as: 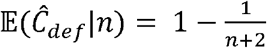. We use this alternative estimate for *Ĉ_def_* when *n* ≤ 5 (cut-off selected via accuracy tests on simulated samples), all the observations are singletons (*f*_1_ = *n*), or if there are no singletons or doubletons (*f*_1_ + *f*_2_ = 0).

### Latent-Trait Model

Species do not interact at random with each other – similar species tend to interact with a similar set of interaction partners. It should be possible to use this information to guide our expectations of missing links through a trait-based framework (Morales-Castilla et al. 2015, Bartomeus et al. 2016, Laigle et al. 2018). These similarities can be in directly-measured traits, such as corolla and tongue length (Vizentin-Bugoni et al. 2014, Klumpers et al. 2019) or through inferred ‘latent’ traits that are not directly measurable. Latent traits can be identified whether the community is structured by phylogeny, microhabitat or physiological differences. A survey of published networks found that the number of latent traits required may be quite low (Eklöf et al. 2013).

Intraspecific trait variation caused by spatial, temporal or ontogenetic differences can pose considerable challenges (González-Varo and Traveset 2016). Nonetheless, such approaches have had successes and a suite of approaches has been developed including matching models (Rohr et al. 2016), Dirichlet-multinomial regression (Crea et al. 2016), matrix factorisation (Seo and Hutchinson 2018) and fourth-corner analyses (Spitz et al. 2014). While it is possible to fit any number of latent traits, here we explore use of a simple single latent trait model to reduce the potential of overfitting (Rohr et al. 2016).

In this model, each focal layer species *i* is assigned a trait value *m_i_* and each partner-layer species *j* a value *m_j_*. The probability of the interaction existing between each *i* and *j* is determined from the difference in trait values through a logistic model:

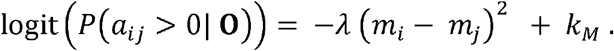

where *λ, k_M_* and the vector of trait parameters **m** are found through maximum likelihood optimisation (details given in SI 3). We constrain *λ* to be positive, such that greater trait differences between potential interactors reduce the probability of an interaction. *k_M_* is an intercept parameter to capture the average probability of an interaction between two species with perfectly matching traits. We penalise strongly divergent traits by introducing a Cauchy-distributed prior with mean 0 on **m**.

Multiple trait distributions may have very similar likelihoods, especially with several specialised interactions or where species bridge network components. To account for this, we optimise 10 differently initialised models and average the predictions of the five models with the greatest likelihood. This model averaging approach is conceptually similar to, but computationally far cheaper than, sampling from posterior distributions of trait vectors.

### Degree Model

It is readily observable that some species have many interaction partners while other species interact with very few. This manifests itself in skewed degree distributions within ecological networks (Jordano et al. 2003). All else being equal, it would be less surprising to discover a missed interaction involving a known generalist species than one with few observed interactions. Many bipartite interaction networks, especially mutualistic networks, are observed to be ‘nested’ - the range of specialists is a subset of the generalists (Bascompte et al. 2003, Ulrich et al. 2009). Simple degree-models can fit binary nestedness and preferential patterns well. Although nestedness can be generated by a variety of processes (Song et al. 2017) and patterns observable in binary networks may not be reflected in the quantitative networks (Staniczenko et al. 2013), our principle objective here is to identify missing links.

To fit a degree model, we assign each species a connectance term, *c*, and determine the probability of an interaction between focal species *i* and interactor *j* as:

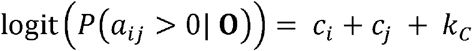

where *k_c_* is a constant intercept term and all parameters are found by maximum likelihood. When both species have high degrees, their connectance parameters will be high, and the probability of those species interacting will be high.

### Matching – Centrality Model

The matching – centrality approach (Rohr et al. 2016, 2018) combines a latent-trait and a degree model and is fit in a similar manner. This dual approach has been shown to have a high capacity to fit tightly to diverse network structures (Rohr et al. 2016). The probability of each interaction existing is found by:

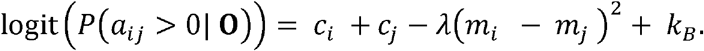

As in the latent-trait model, *λ* is defined to be positive, *k_B_* is a constant intercept term and all parameters are found by maximum likelihood. We again apply a weak Cauchy prior centred at 0 onto the latent trait terms **m** and fit 10 models, averaging predictions of the best five.

### Block Model

Ecological networks, especially antagonistic networks, frequently show comparatively discrete compartments or modularisation, where subsets of species interact more strongly within their group than with outsiders (Olesen et al. 2007, 2008, Allesina and Pascual 2009, Schleuning et al. 2014). This may reflect fundamental incompatibilities in physiological traits, temporal mismatch (such as nocturnal/diurnal partitioning) or spatial segregation.

This grouping can be represented by stochastic block models (SBMs), which have been shown to perform well on ecological data in comparison to other clustering algorithms (Leger et al. 2015) and are used increasingly in ecology (Sander et al. 2015, Kéfi et al. 2016). In SBMs, each species is assigned to a group, *G_x_*, in a defined set: *x* ∈ {1,…, *g*}. The probability of interaction between two species is determined based on their group membership:

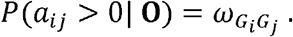

The elements of *ω* are the between-group interaction probabilities and are directly specified as the fraction of observed interactions between each of the groups. We find optimal group assignations and fit the model using a degree-corrected bipartite-SBM specific algorithm (Larremore *et al.* 2014, detailed in SI 3). To capture inherent indeterminacy in underlying group structure, we fit 10 differently-initialised models, and use the average prediction of the five models with the highest likelihood.

### Combinations of Models

Individually the above models each capture discrete pieces of information about the identity of missing links. However, the structure of ecological networks is the product of many separate drivers. To capture this diversity, we combine the predictions of multiple models into ensembles. We test combining the matching-centrality model with the block model, and each of the ‘structural’ models with the coverage-deficit model.

We test two ensembling approaches, multiplication and averaging. Multiplying the relative probabilities assigned to each putative missing link, *P_ens_*(*a_ij_* > 0) ~ *P*_1_(*a_ij_* > 0) × *P*_2_(*a_ij_* > 0), emphasises the extreme probabilities of the constituent models. Averaging the relative probabilities, *P_ens_*(*a_ij_* > 0) ~ *P*_1_(*a_ij_* > 0) + *P*_2_(*a_ij_* > 0), highlights possible interactions that are consistently identified by multiple models. Before combining models we standardise each set of probabilities assigned to unobserved interactions to sum to 1.

#### Datasets

Testing the efficacy of extrapolations requires knowledge of the ‘true’ network. We take two complimentary approaches to defining our ‘true’ networks. First, we generate a set of simulated networks that vary over a wide range of network properties. Second, we use a large and diverse set of empirical networks from literature sources. In both cases ‘observed’ networks were generated by taking draws from a multinomial distribution parameterised by the ‘true’ interaction frequency. This replicates ecological sampling processes where the most frequent interactions are highly likely to have been observed and the rarest missed.

We initially generated 2000 simulated networks from a probabilistic two-trait niche model (modified from Fründ *et al.* 2016) described in full in SI 1. Key network properties were calculated using the *bipartite* R package (Dormann et al. 2008): connectance, weighted nestedness (Galeano et al. 2009), cluster coefficient (Watts and Strogatz 1998), distribution of interaction frequency (Shannon diversity, Bersier *et al.* 2002) and network level specificity H2’ (Blüthgen et al. 2006). Distributions of these metrics and the (low) correlations between them are depicted in SI 1.

Our objective was to generate interaction matrices that represent a wide range of ecologically plausible bipartite networks. However, models used to generate plausible interaction networks have many similarities to the predictive models described above. This is for a good reason – models are chosen because they are thought to represent ecological networks parsimoniously. To reduce circularity, our generative model is considerably more complex than our predictive models and we examine how predictive model performance changes in response to network properties.

Using the ‘true’ networks we took 300-2000 samples per network to generate our ‘observed’ networks. We discarded occasional cases where all true interactions were observed. This resulted in a low mean true sample coverage deficit of 2.4%, corresponding to an average network completeness of 64% due to skewed interaction strength distributions. Our ‘observed’ networks therefore represent reasonably ‘well-sampled’ systems that are nonetheless missing a large proportion of the interactions present.

We collated a diverse set of 113 empirical networks representing antagonistic, mutualistic and commensalistic interaction types (SI 2). We collated quantitative single-class bipartite networks from the *Web of Life* (www.web-of-life.es) repository with 30-200 species and 40-431 observed interactions. This included 48 plant-pollinator (19 sources), 23 mammal-flea (one source, but across a wide geographic range), 2 plant-ant mutualism (2 sources) and 10 seed disperser (from 7 sources) networks. We supplemented this with 25 mammal-dung beetle interaction networks (Frank *et al.* 2018, selecting those with ≥30 total species, ≥40 observed interactions, and ≥5 mammal species) and 5 host-parasitoid networks (Tylianakis et al. 2007).

These empirical networks are not exhaustively sampled - the Chao1 estimator indicates that the observed interactions represent on average only 76% of the underlying network (SI 2). Nonetheless, we took the empirical interaction frequency to be the ‘true’ network to define a multinomial distribution. We subsampled each empirical network 20 times, each time taking a different fraction between 0.2 and 1 of the original sample size, taken to be the sum of the observations. We excluded cases where the observed web included more than 95% of the ‘true’ interactions.

#### Model Testing

Using each model or combination of models we assign each unobserved interaction a relative probability of being a missing link. We assume that all observations are true positives (i.e. there have been no misidentifications) although in principle similar methods could be used to identify candidates for re-evaluation. Where a species in the ‘true’ network was not included in the ‘observed’ network, we did not attempt to predict its interactions and reduced the size of the ‘true’ network correspondingly for all purposes.

### Identifying missing links

We assess performance at identifying missing links with the area under receiver operating characteristic curve (AUC) metric to assess the information content of a signal, using the pROC R package (Robin et al. 2011). AUC can be considered the chance that a ‘false negative’ (missing link) was assigned a higher relative probability than a ‘true negative’. A value of 0.5 indicates no useful signal while a value of 1 indicates perfect discrimination. To determine if *P*(*a_ij_* > 0) is reflective of true relative interaction frequency, we calculated the mean Spearman’s rank-correlation with the ‘true’ relative interaction strengths of the unobserved interactions for each model.

### Improving estimates of network metrics

We used the *estimateR* function in the *vegan* R package (Oksanen et al. 2018) to estimate the number of missing links based on the total distribution of observed interaction counts using the bias-corrected Chao1 index. We then augmented the observed interaction networks by adding the estimated number of the most probable missing links to the network based on the average of the block, matching-centrality and coverage-deficit models. Each selected missing link was taken to be ‘observed’ once, and network-level metrics calculated for the observed, augmented and true networks. We tested three qualitative network metrics (connectance, nestedness and niche overlap) identified as being susceptible to misestimation in under-sampled networks (Fründ et al. 2016), alongside two metrics considered to be comparatively resistant to under-sampling (H2’ and weighted NODF nestedness).

## Results

### Identifying missing links - Simulated networks

In general, the network models showed impressive performance at identifying likely missing links. The best-performing model overall was the block model, closely followed by the other three network models (Fig. 2a). The coverage deficit model lagged behind the structural models, but was more informative than simply ranking focal species by sample size. Combining the predictions of multiple models tended to improve performance relative to individual models (Table 1). The overall best-performing model averaged the predictions of the block model, the matching-centrality model and the coverage-deficit model. However, the marginal improvements from combining models were often small.

**Figure 2.**
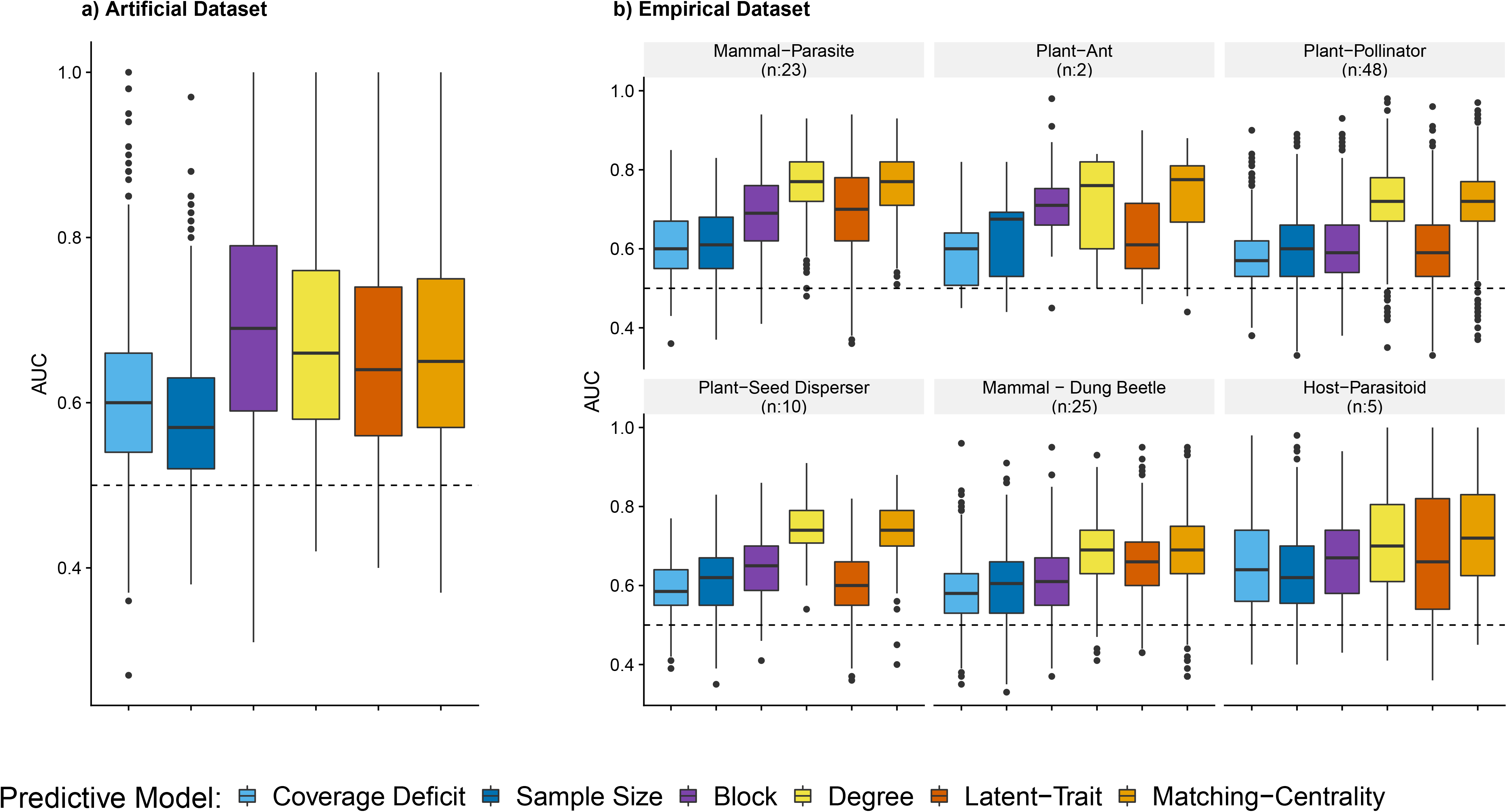
**a)** Model performance across 1985 simulated networks. Mean differences between the statistical models were small, although given the large sample size all pairwise differences were statistically significant (Holm-corrected t-tests, paired by network ID, p<0.0001). **b)** Model performance on the empirical data sets, subdivided by network class. Each empirical network was subsampled 20 times to different fractions of the original sample size. Models perform differently on different classes of data (Model comparison, accounting for dataset ID as a random effect, shows model:class interaction term is significant: p< 0.0001, *n_obs_* = 12672). In all cases boxes show first and third quartiles.

**Table 1.** Mean AUC values using different predictive models or combinations of models across the whole simulated (n=1985) and empirical (n=113, 2112 subsamples) network sets. Within each dataset, results are shown with just the network-structure based model (left column), ensembled with the coverage-deficit model, by multiplying the probabilities (centre column) or by averaging the assigned probabilities (right column). Incorporating coverage deficit tends to lead to modest improvements in the simulated networks but not in the empirical networks. Sets of results found not to differ significantly (Holm-correct t-tests, paired by network ID) are indicated with grouping letters, applied separately for simulated and empirical results.

| Model | Simulated Dataset<br>Accuracy Score<br>(AUC) |  |  | Empirical Dataset<br>Accuracy Score<br>(AUC) |  |  |
| --- | --- | --- | --- | --- | --- | --- |
|  | Structural<br>Model-Only | Ensembled with<br>Coverage-Deficit model |  | Structural<br>Model-Only | Ensembled with<br>Coverage-Deficit model |  |
|  |  | (×) | (+) |  | (×) | (+) |
| *Focal Layer Sampling* |  |  |  |  |  |  |
| Coverage-Deficit | 0.600 |  |  | 0.589<sup>abc</sup> |  |  |
| Observation Count | 0.580 |  |  | 0.603<sup>de</sup> |  |  |
| *Network Models* |  |  |  |  |  |  |
| Latent-Trait | 0.660<sup>a</sup> | 0.649 | 0.654<sup>a</sup> | 0.631<sup>f</sup> | 0.588<sup>abc</sup> | 0.590<sup>abc</sup> |
| Degree | 0.676 | 0.708<sup>efg</sup> | 0.683<sup>b</sup> | 0.721<sup>i</sup> | 0.651<sup>h</sup> | 0.663 |
| Matching-Centrality | 0.671 | 0.702<sup>de</sup> | 0.684<sup>b</sup> | 0.718<sup>i</sup> | 0.642<sup>g</sup> | 0.658 |
| Block | 0.691<sup>bcd</sup> | 0.689<sup>bc</sup> | 0.695<sup>cd</sup> | 0.630<sup>f</sup> | 0.584<sup>a</sup> | 0.590<sup>c</sup> |
| *Combined Models* |  |  |  |  |  |  |
| Block and Matching-Centrality (Multiplied) | 0.702<sup>de</sup> | 0.705<sup>ef</sup> | 0.711<sup>f</sup> | 0.647<sup>gh</sup> | 0.594<sup>c</sup> | 0.604<sup>de</sup> |
| Block and Matching-Centrality (Averaged) | 0.705<sup>ef</sup> | 0.709<sup>f</sup> | 0.713<sup>g</sup> | 0.648<sup>gh</sup> | 0.600<sup>d</sup> | 0.603<sup>d</sup> |

Model performance and the relative hierarchy of models was affected by the underlying true network structure (Fig. 3). On average, missing links were identified best in nested, unclustered networks where interaction frequencies were more evenly distributed. The coverage-deficit and sample size models were the least responsive to network structure, since our simulated network generator did not result in a pronounced abundance-generality correlation. Average model performance did not change appreciably with increased sample size, overall interaction coverage deficit or samples per host (Fig. S4). However, when a greater fraction of interactions had been observed the structural models increased their performance, while performance of the coverage deficit model remained unchanged (Fig. S4).

**Figure 3.**
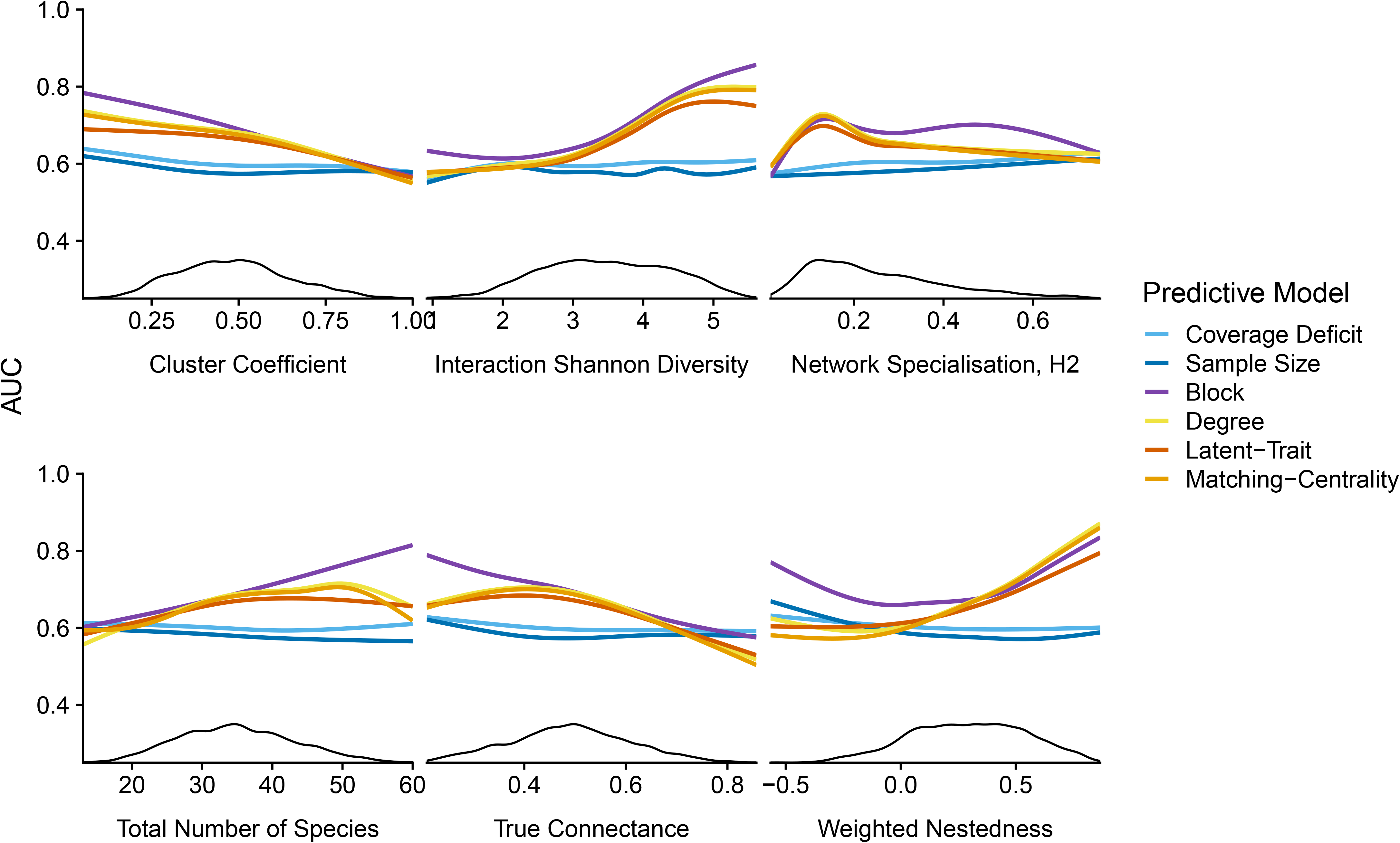
Response of model performance to ‘true’ network structure. Smoothed line shows a GAM fit for each predictive model. Lower black line shows the local density distribution of that property across the simulated test dataset.

### Identifying missing links - Empirical networks

The performance of the different predictive models varied considerably between the different categories of empirical network (Fig. 2b). As expected, the degree model performed best for mutualistic networks, and the latent-trait model for antagonistic networks. In all six network categories the matching-centrality model had the best overall performance. The ‘true network’ properties (Fig. S3) or the degree of sampling rarefaction (Fig. S4) had relatively little effect on predictive performance, except for networks that show ‘anti-nestedness’. In these cases, the predictive capacity of the degree model was low, although the matching-centrality model maintained its advantage. The coverage-deficit model was very poor at predicting missing links in the empirical datasets and combining it with the network-structure models led to a *reduction* in overall predictive power (Table 1).

### Identifying interaction frequency of missing links

Correlations between probabilities assigned to missing links *P*(*a_ij_* > 0|**O**) were related to their ‘true’ interaction frequency. The relationship was stronger in the simulated dataset than the empirical dataset (Table S2). Of the structural models, the degree model showed the strongest positive correlation (*ρ_Sim_* = 0.51, *p_Emp_* = 0.15), while the block model showed the weakest (*ρ_Sim_* = 0.28, *ρ_Emp_* = 0.06). The coverage-deficit model was negatively correlated with true frequency (*ρ_Sim_* = −0.35, *ρ_Emp_* = −0.05) i.e. missing links successfully identified via this model tended to be infrequent.

### Improving estimates of network metrics

Integrating inferred interactions into observed networks could improve estimated of key metrics. The ‘augmented’ networks showed increased correlation, reduced bias and reduced error in qualitative metrics, compared to the original observed networks (Fig. 4a). The quantitative network metrics (H2’and weighted NODF, Fig. 4b) were already well-estimated by the under-sampled network, and the inclusion of additional interactions made little difference. The Chao1 estimator for the number of missing interactions appeared to perform well for the empirical networks but consistently underestimated the true interaction count and coverage deficit in the simulated data (Fig. S5).

**Figure 4.**
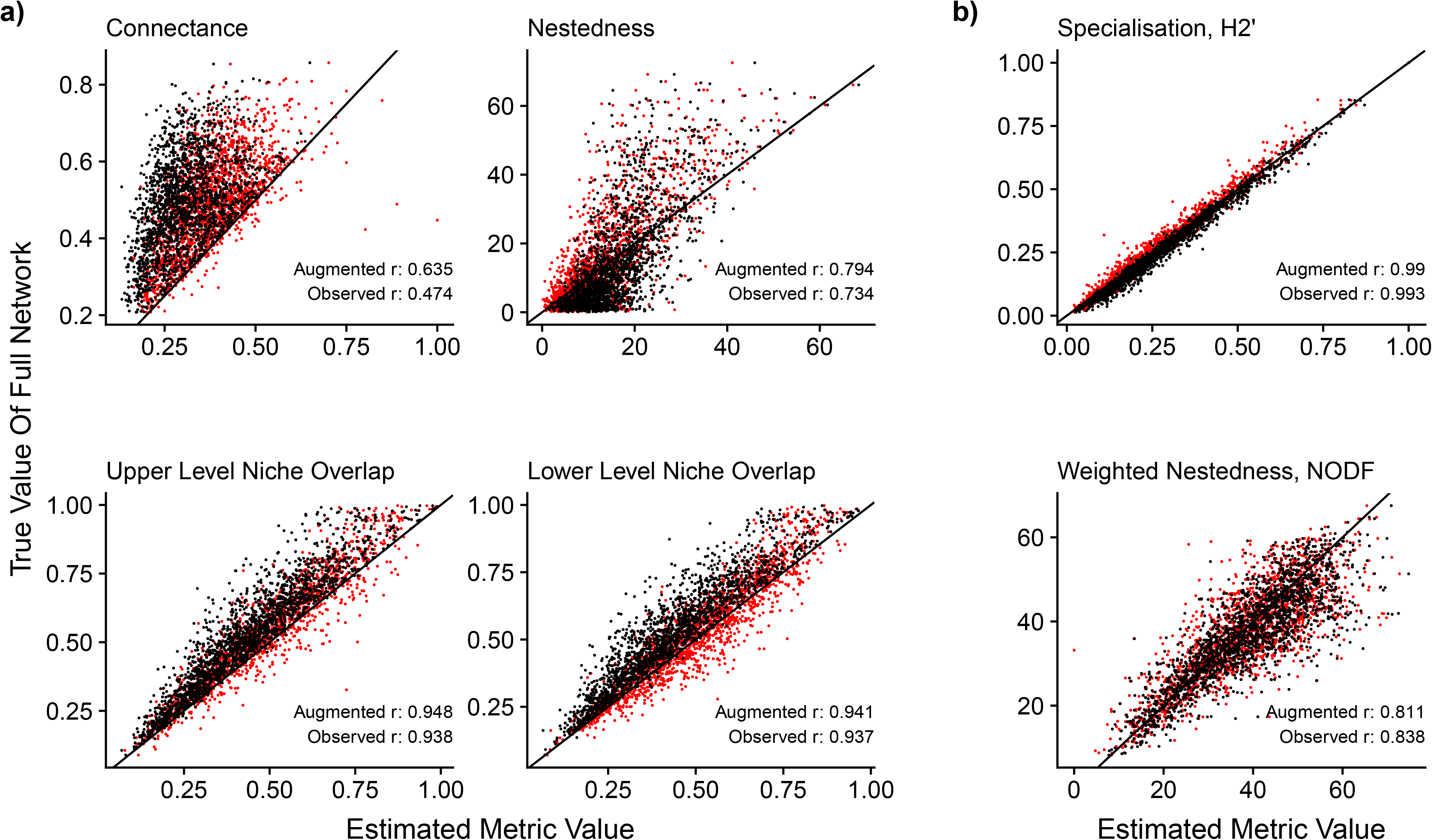
Relationship between network metrics calculated on the observed (black) and ‘augmented’ (red) interaction networks to the true network value in the simulated networks. The line shows the 1:1 relationship. Pearson’s correlation coefficient (r) is shown for each metric. Qualitative metrics (a) show notable improvements in correlation with the true values when augmented, while quantitative metrics (b) are largely unchanged.

## Discussion

We have shown that is possible to infer missing links in range of ecological networks with a reasonable degree of accuracy and this approach can improve estimates of network metrics. The challenge now is to put these tools into practice to leverage additional ecological insight.

### Choosing between predictive models

Ideally, a model used to infer missed interactions would capture the true ecological drivers, but this is not essential for all purposes. Given the diversity of ecological networks, there will never be a single ‘best’ model and victors in comparisons will depend on the data set. In our simulated data our block model appears to perform best. In our empirical datasets, the matching-centrality model performed consistently well (as found by Rohr *et al.* 2016). However, we have reason to distrust the empirical datasets (discussed below). The pronounced, likely artefactual, abundance-generality relationship will favour degree-models.

Nonetheless, splitting hairs over the best-performing model is not necessarily a productive route. Networks are structured by multiple processes and in our simulated data sets the very best performance comes from combining different models, which identify different missing interactions. Structure-based models pick out the more frequent interactions while coverage deficit models highlight comparatively infrequent interactions that would be the hardest to determine through further undirected sampling. Future progress will come from operationalising estimated missing links, rather than from further incremental model refinements.

### Distinctions between empirical and simulated networks

The empirical and simulated datasets overlapped substantially in key network metrics. The main relevant difference between these datasets is the stronger correlation in empirical networks between species’ marginal totals and generality (mean Spearman’s *ρ_Sim_* = 0.43, *ρ_Emp_* = 0.76). This can account for the poor performance of the coverage-deficit model in these cases. Disentangling the extent to which apparent specialism of rarely observed species is a sampling artefact is challenging (Blüthgen et al. 2006), since there are ecological explanations for such a relationship (Fort et al. 2016, Dormann et al. 2017, Simmons et al. 2019). For our purposes, this likely bias in the structure of empirical ecological networks has the consequence that predictive models may identify missing links introduced both by our subsampling procedure, and due to gaps in the original dataset. We therefore place more weight on the results from the simulated data, while noting that, despite the obstacles, the structural models are still able to perform reasonably well on the sparse empirical data.

### Estimating the quality of sampling and the number of missing interactions

Sample coverage estimators offer promise for estimating the number of missed interactions (Chacoff et al. 2012, Traveset et al. 2015, Jordano 2016, MacGregor et al. 2017). The distinction between sample-coverage and network completeness is worth emphasising. Apparently high values of sample coverage alone should not be used as a simple proxy for ‘network completeness’, although they are related to the reliability of certain quantitative network metrics (Henriksen et al. 2019). We found that the whole-network-level Chao1 estimate of missed interactions was strongly correlated with the ‘true’ number of missing links in our simulated networks, but notably underestimated this number, a pattern also observed by Fründ *et al.* (2016). Although there appeared to be no systematic bias in estimated network completeness in the empirical datasets, these ‘true’ networks are themselves highly incomplete, suggesting an overall underestimation also in these cases. Our focus in this paper is on the location rather than the number of missing links. Future work should investigate how long tails of infrequent interactions relate to this underestimation, and consider whether species rank abundance distribution estimators (Chao et al. 2014) can be usefully generalised to assign a frequency distribution to estimated missing links.

### The way forward

Most of the models used here, and indeed our fitting approaches, likely have further room for refinement, and there are many further classes of predictive models that we did not test, such as nearest neighbour algorithms (Desjardins-Proulx et al. 2017). Furthermore, we based our models entirely on the information contained in the network itself. In most cases, ecologists constructing a network have access to (sometimes considerable) additional information and expertise on the study system. Introducing pre-existing knowledge about likely or forbidden interactions, or pooled knowledge from multiple sample sites offers considerable opportunities to maximise the information in a network (Gray et al. 2015, Pomeranz et al. 2018). External information would be particularly useful in separating ‘forbidden links’ (Olesen et al. 2011) that never occur due to fundamental incompatibilities from those that occur elsewhere but not within those spatio-temporal boundaries. For instance, species may not be observed to interact because of a fundamental mismatch in body-sizes (a forbidden link) or because they interact at night but data were gathered only during the day.

Because the naïve models used here do not use external information, or the judgment of the ecologists constructing the network, the quality of the estimation very much represents a lower bound of what is possible with the approach in practice. As a result of the underlying idiosyncrasies of ecological systems, often euphemistically labelled ‘species identity effects’, there will always be an upper limit to the predictive capacity of any statistical model. We suggest that, rather than developing new models to gain marginal improvements in predictive capacity, a more productive focus would be to develop frameworks exploiting this information to test the robustness of conclusions derived from ecological networks.

Complete network inventories are not a realistic or necessarily useful goal (Jordano 2016). Enormous sample sizes and strictly deliminated boundaries are required to exhaustively sample all interactions (Martinez et al. 1999). There is undoubtedly a long tail of extremely infrequent interactions that will never be observed in a realistic sampling regime. Furthermore, there is rarely a clear line defining network boundaries (Poisot et al. 2015). Ecological networks are a constantly moving target (Rasmussen et al. 2013, CaraDonna et al. 2017) and there may be little value in including certain rare, occasional visitors and unusual interactions. It can be challenging to determine if infrequent interactions are simply a curiosity (e.g. Mercer 1966) or have significant consequences (e.g. Dudley et al. 2016). The utility of increasing the accuracy with which one type of interaction is understood will plateau as the impact of other interaction types and ecological guilds predominate (Lafferty et al. 2008). Despite the challenges, we see two principle uses for inferred missing links.

First, inferring missing links will direct further sampling where the goal is a descriptive network. In many cases the topography of the network is of principle interest, given the potential for interaction strengths to vary through time and in response to perturbation. For example, when tracing the relationships of species that may be the target for eradication measures, such as disease-vector mosquitoes (Collins et al. 2019), a critical first step is an inventory of interactions in the community. Since in most cases investment in sampling is a finite resource, directing sampling towards high-likelihood interactions could more efficiently increase the overall network completeness. However, we caution that directed sampling will introduce biases in the generated network (Gibson et al. 2011, Falcão et al. 2016) and may hinder cross-community comparability (Poisot et al. 2012a). In certain cases, independently-sampled networks (even if less complete) may be more useful than comparisons between ‘augmented’ networks. The use of information from other networks may lead them to appear more homogenous than they really are and lead to underestimates of interaction β-diversity through time and space (Poisot et al. 2015, Graham and Weinstein 2018).

Second, the identification of probable missing links can improve tests of the reliability of conclusions drawn from networks, potentially through explicitly probabilistic networks (Poisot et al. 2016a). Statistical resampling and rarefaction procedures that can only work backwards or by adding random interactions may both introduce biases. While particular quantitative metrics are relatively robust to under-sampling (Blüthgen et al. 2008, Fründ et al. 2016, Henriksen et al. 2019), these metrics may not align with the goals of the study. If the conclusions are robust to the inclusion of probable missing links, then confidence in the results would be increased.

Conversely, if including most-likely missing links alters conclusions, this would signal that these conclusions are unlikely to be robust. In these cases, care should be taken when selecting a predictive model to avoid a clash with the properties under investigation.

### Conclusion

The identification of likely missing links is a powerful tool to include in the arsenal of network ecologists. We have shown that models to identify missing links can perform well across a wide range of network types. Importantly, inferential models need not be technically challenging to implement – for example the *cassandRa* package allows all the models discussed here to be fit with a single function call. We hope that such approaches become more widely used in guiding ecological sampling and testing the robustness of conclusions drawn from bipartite networks.

## Acknowledgments

Thanks to the CERO group for comments on this project, and two anonymous reviewers for their constructive comments. Funding was provided by NERC Standard Grant NE/N010221/1.

## Data accessibility

R functions to run the predictive models are publicly available in the R package *cassandRa* on CRAN. All code and necessary data to generate simulated datasets and carry out analyses are available on GitHub (github.com/jcdterry/PredictingMissingLinks). Should the manuscript be accepted, this will also be archived in a public repository and the data doi included.

## Statement of Authorship

JCDT initiated the research, in discussion with OTL. JCDT conducted all analyses and wrote the first draft of the manuscript. Both authors contributed to revisions.

## Notes

#### Summary of Updates

Moving more of the Methods from SI to main text and improving presentation of results.

