## Supplementary Information for "Finding missing links in interaction networks"

**4 Supplementary Results** **8**

**5 Original Empricial Network Sources** **11**

#### List of Tables

- 1 Network generation parameters used to generate simulated dataset. 2
- 2 Spearman's rank correlations between the missing link probability with each model and the 'true' relative interaction frequency. . . 8

### 1 Simulated Communities

#### 1.1 Generation of networks

Networks were generated with the `make_true_and_sample_web()` function in the `cassandRa` R package (Available on CRAN). This function takes a random number generator seed, the dimension of the desired network, a number of sample observations and five network parameters.

We made 2000 communities, although in 25 cases the sampled community already contained all the interactions so were discarded, leaving a total dataset of 1985. The parameters were all independently drawn form distributions listed in Table S1.

| Parameter | Meaning | Distribution |
|---|---|---|
| $n_{focal}$ | Number of species in focal level | Integer uniform distribution: 15-30 |
| $n_{interactor}$ | Number of interaction partner species | Integer uniform distribution: 15-30 |
| $s$ | Specialisation parameter | One of: 0.5,1,2,5,10, 50 |
| $\sigma_{abund}$ | Log-Abundance standard deviation | Uniform distribution: 0.5 - 3 |
| $\tau$ | 'trait' and 'nestedness' balance | Uniform distribution: 0.1 - 0.7 |
| $\theta$ | Number of sample observations to draw | Integer uniform distribution: 300-2000 |
| $\gamma$ | Target connectance of true network | Uniform distribution: 0.2 - 0.5 |

Table 1: Network generation parameters used to generate simulated dataset.

#### 1.2 Network Generating Model

The aim of this procedure is to generate networks that span the full range of ecological bipartite interaction matrices. The distribution of these networks is not therefore intended to match the distribution of known empirical networks, since these will be skewed by choices of which networks to target.

A species interaction preference matrix ($P_{ij}$) was determined using a two trait model based on that used by Fründ *et al.* (2016). Each species was assigned two trait values drawn from a uniform distribution. The Euclidean trait distance between host $i$ and interactor $j$ was then calculated, placing twice the weight on the $\alpha$ trait:

$$T_{ij} = \sqrt{(\alpha_i - \alpha_j)^2 + \left(\frac{\beta_i - \beta_j}{2}\right)^2}$$

Trait difference were translated into a preference using a Gaussian model and the specialisation parameter $s$:

$$P_{ij} = \exp(-s^2 T_{ij}^2).$$

Where $s$ is higher, the preference drops away faster as the difference in trait space increases. Because species are unevenly distributed in trait-space, this results in a somewhat uneven degree distribution.

A specialisation matrix ($N_{ij}$) is then generated. First each focal species is independently assigned a degree term $c$ drawn from a Poisson distribution, parameterised based on the desired target connectance and the number of species in the other bipartite layer:

$$c_{focal_i} \sim Pois(\lambda = \gamma n_{interactor})$$

$$c_{int_j} \sim Pois(\lambda = \gamma n_{focal})$$

These vectors of species level parameters are then multiplied to generate the nestedness matrix:

$$N_{ij} = c_{focal_i} \times c_{int_j}$$

The two interaction frequency matrices are then each scaled by their root square mean and combined, weighted by the $\tau$ parameter:

$$M_{ij} = P^*_{ij}(\tau) + N^*_{ij}(1 - \tau)$$

A degree of noise is added to this matrix by multiplying each element by an independently drawn value from a uniform distribution between 0.9 and 1.1 to generate $M'_{ij}$, an overall matrix of per-individual interaction rates.

The target number of interactions is calculated as $\psi = n_{focal} \times n_{interactor} \times \gamma$. The largest $\psi$ values of $M'_{ij}$ are retained while the remainder are set to 0. This introduces forbidden links without reference to the relative abundance of either species.

The relative abundance $x_i$ of each species is assigned by taking values that conform to a log-normal distribution with mean=5 and sd=$\sigma_{abund}$, following the approach of Fründ *et al.* (2016). From this the true relative interaction frequency matrix $\mathbf{A}_{ij}$ is calculated as: $\mathbf{A}_{ij} = x_i x_j M'_{ij}$ .

Elements of the observed interaction matrix were drawn from a multinomial model with $\theta$ draws. In cases where there were no observations of any species, those rows or columns of the matrices were removed. Hence, we only attempt to infer interactions between the observed set of species for which we have some information about. We exclude cases where there are no missing links to be found because all interactions had been observed at least once.

#### 1.3 Properties of Simulated Datasets

The distribution of several key network properties are shown by the density plot in main text Figure 3. The rank correlation between the set parameters (first four row/columns) and network metrics of the generated webs are shown in the correlation plot Figure S1.

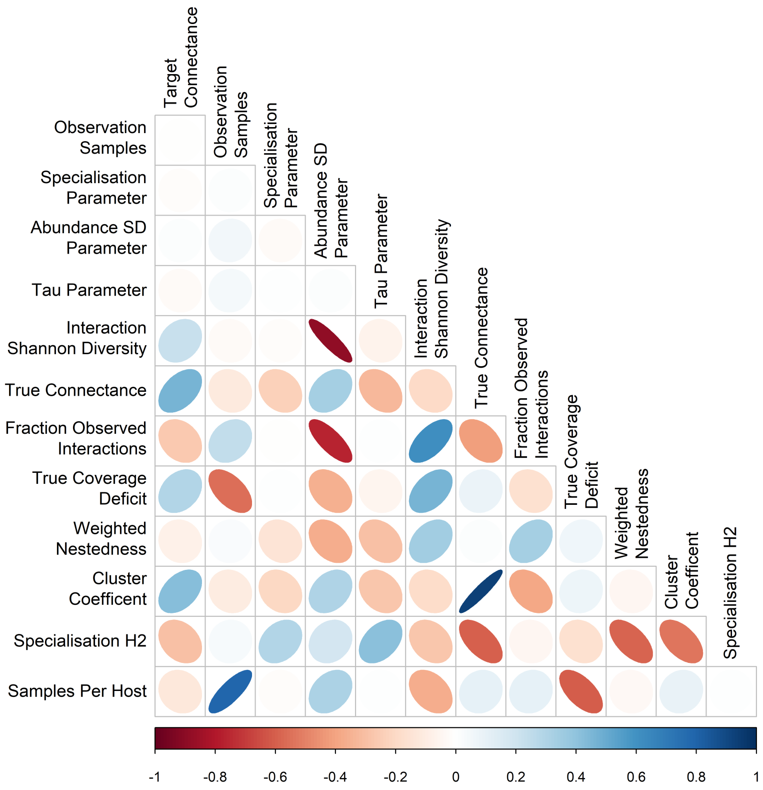

Figure S1: Spearman's Rank Correlations between metrics of the simulated networks. First 4 rows are set parameters, and hence uncorrelated with each other. Tighter, darker ellipsis indicate stronger correlations. Direction and colour of ellipses indicate sign of correlation.

### 2 Empirical Datasets

Original data references given in the original collations of datasets (www.web-of-life.es) and Frank *et al.* (2018) are listed in SI 4. Published networks were treated as raw observation counts in all cases. In the one case of non-integer interaction frequencies, the values were rounded.

Sample coverage of the empirical networks was estimated using the Chao1 estimator and the number of estimated unobserved interactions was calculated by comparing the estimated interaction count from the `estimateR` function in `vegan` package with the observed fraction.

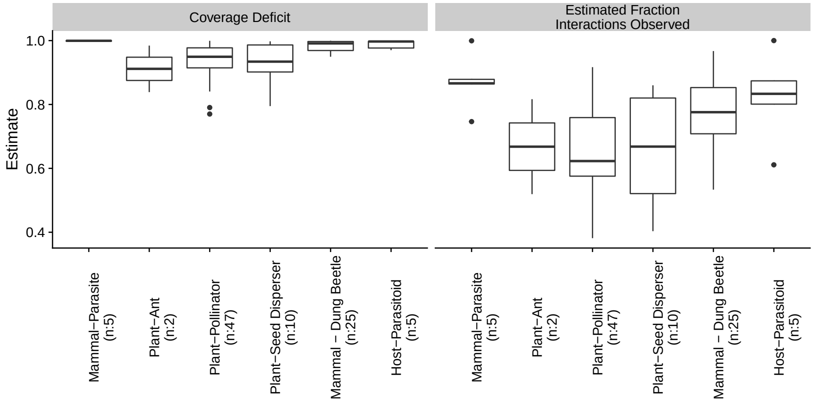

Figure S2: Estimated coverage deficit and estimated fraction interactions observed (network completeness) of the empirical networks collated. Only 5 of the mammal-parasite networks had sufficient singleton interactions to meaningfully estimate interactions observed.

### 3 Predictive Models

All model fitting was conducted using the `cassandRa` R package (Terry 2019) version 0.1.0 available on CRAN.

#### 3.1 Optimisation Procedures for Trait, Degree and Matching-Centrality models

Maximum likelihood estimates of best-fitting parameters for each of the models were found using the optim function and the Nelder-Mead method. Each fit was run up to 10'000 steps. Each model was fit 10 times and the predictions from the best (highest maximum likelihood) models were averaged. This procedure gave improved performance at finding well-fitting models compared to extending the maximum number of iterations and had an averaging effect when there were multiple near-equivalent models

Starting parameters were chosen as follows:

- Degree terms ($c_i$): drawn from Gaussian distributions (mean 0 and sd 1).
- Latent trait terms ($m_i$): column and row sums of a CCA ordination are used to find likely start points
- Constant terms ($k$): drawn from a uniform distribution 0:1
- Trait-weighting term ($\theta$) : drawn from a uniform distribution 0:1

#### 3.2 Details of the Stochastic Block Model

For the SBM, each species is assigned to a group, $G_x$, in a defined set: $x \in \{1, \ldots, g\}$. The probability of interaction between two species is defined based on their group membership:

$$P(a_{ij} > 0) = \omega_{G_i G_j}.$$

To parameterise the model, the $\omega$ matrix is directly specified as the fraction of observed interactions between each of the groups. Specifying the number of groups and assigning species to groups is more challenging. With an increased number of groups, $g$, the quality of the fit will improve, up to the point where each species assigned its own group and the model fits perfectly. The size of the matrix $\omega$, and hence the number of fitted interaction probability parameters is $g^2$. While methods exist to determine the most parsimonious $g$ to detect clustering, here we set $g$ to be the square root of the number of species in the smaller bipartite layer, rounding down to a minimum of two.

We find optimal group assignations using a degree-corrected bipartite-SBM specific algorithm (Larremore *et al.* 2014). This approach ensures that the model does not need to learn the underlying bipartite structure and the proclivity to group highly connected species is tempered. To capture uncertainty in underlying group structure, we fit 10 differently-initialised models, and use the average prediction of the best five models.

Because group-assignments in an SBM are discrete, gradient-based optimisation methods (such as those employed by the `optim` R function) are not applicable. We follow the stochastic optimisation algorithm of (Larremore *et* *al.* 2014) to find good assignations. We again fit 10 models and average the predictions of the best 5 in order to avoid inclusion of poor local optima and to incorporate uncertainty in group assignation.

Given a particular set of group assignations, the likelihood of the model is calculated a group-combination at a time. The likelihood contributions of each group r interacting with each group s is given by:

$$\mathcal{L} = \sum_{rs} \sum_{i \in G_r} \sum_{j \in G_s} \mathbf{A}_{ij} \ln \frac{\mathbf{A}_{ij}}{\kappa_i \kappa_j}$$

Where $\mathbf{A}$ is the bipartite adjacency matrix, $\kappa_i = \sum_j \mathbf{A}_{ij}$ is the sum of degrees in the group. Where many interactions are included in some group-pairs and few interactions are included in other group-pairs, the likelihood will be maximised. In contrast if the number of interactions per group-pair is relatively constant then the likelihood will not be as high. Dividing by the degree correction term $\kappa_i \kappa_j$, rather than the number of species in the group, restrains the model from grouping all generalist species together in a group.

The algorithm used to find high-quality group assignations is as follows. Initially, all species are randomly assigned to groups. Then, one at a time, each species is swapped into a different group (at random) and the likelihood of the assignations assessed as described above. The best model of all these swaps is then selected (even if it is worse than the original) and used in the next round of swapping. This is repeated until a stopping criteria is met: no more than a 0.1 log-likelihood improvement in the best model of the last 20 selected compared to the best overall model.

Between-group interaction probabilities $\omega$ are then found as the fraction of interactions between each group that had been observed.

### 4 Supplementary Results

| Model | Simulated | Empirical |
|---:|---|---|
| Coverage Deficit | -0.346 | -0.047 |
| Degree | 0.517 | 0.154 |
| Latent-Trait | 0.335 | 0.061 |
| Matching- Centrality | 0.517 | 0.136 |
| SBM | 0.276 | 0.059 |

Table 2: Spearman's rank correlations between the missing link probability with each model and the 'true' relative interaction frequency.

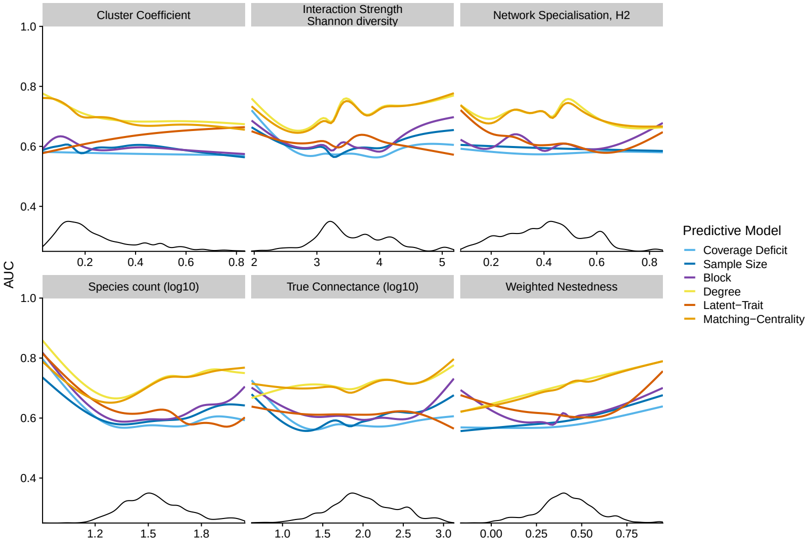

Figure S3: Effect of various network properties on the capacity of different predictive models to identify missing links in the empirical networks. Lines show smooth lines fit with a GAM. Black density plots show distribution of that structural property across the emprical networks.

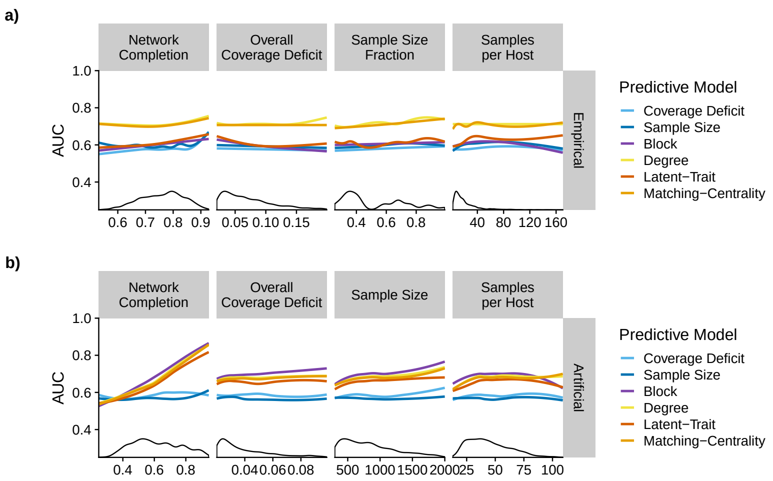

Figure S4: Impact of sampling quality on predictive performance of different predictive models on a) empirical datasets, b) simulated datasets. Lines show smooth lines fit with a GAM. Black density plots show distribution of that structural property across the emprical networks

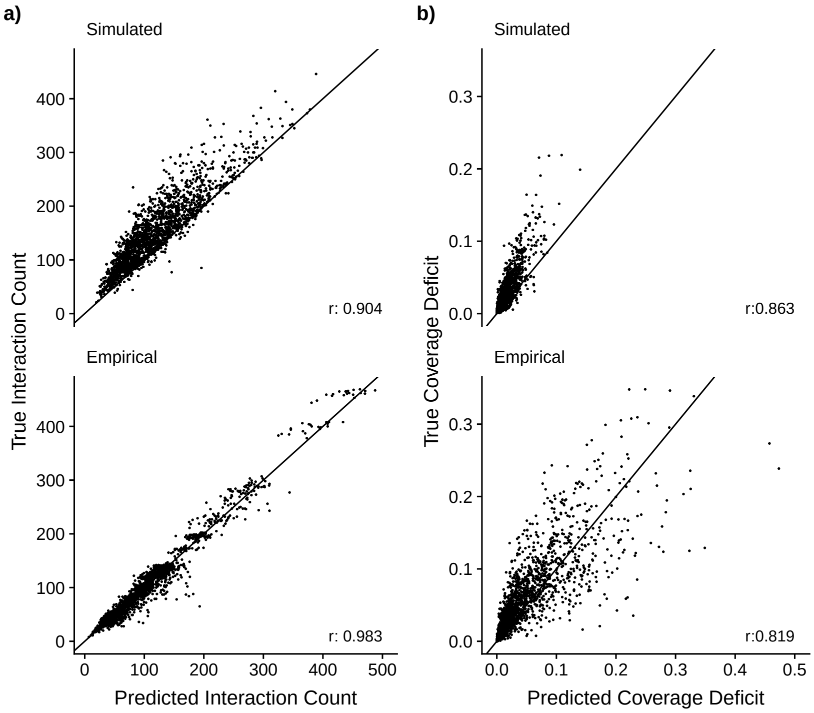

Figure S5: Relationships between a) estimated sample coverage and 'true' sample coverage and b) estimated number of missing interactions and the 'true' number of missed interactions. Results are shown using the bias-corrected Chao1 estimator, the ACE estimator had very similar results. Line shows a 1:1 relationship that an unbiased metric would follow.

### 5 Original Empricial Network Sources

Original data sources for empirical networks, as cited and labelled by the compiling papers, along with network size, basic connectance and location where available. Raw names of networks as provided by those collations are also included.

| ID | Focal Layer Species | Interaction Partners | Connectance | Type of interactions | Reference | Locality of Study | Latitude | Longitude |
|---|---|---|---|---|---|---|---|---|
| A_HP_002 | 42 | 96 | 0.222 | Host-Parasite | Hadfield et al. (2013) The American Naturalist 183:174-187 | Akmolinsk | 51.17 | 71.42 |
| A_HP_003 | 32 | 108 | 0.522 | Host-Parasite | —"— | Altai | 49.00 | 89.00 |
| A_HP_006 | 53 | 123 | 0.208 | Host-Parasite | —"— | Armenia | 40.18 | 44.52 |
| A_HP_009 | 36 | 97 | 0.315 | Host-Parasite | —"— | Dzhungarsky Alatau | 45.00 | 80.00 |
| A_HP_010 | 49 | 88 | 0.158 | Host-Parasite | —"— | East Balkhash | 46.17 | 74.33 |
| A_HP_013 | 33 | 103 | 0.426 | Host-Parasite | —"— | Guriev | 47.26 | 52.06 |
| A_HP_018 | 36 | 127 | 0.403 | Host-Parasite | —"— | Kostroma | 57.77 | 40.93 |
| A_HP_020 | 33 | 105 | 0.386 | Host-Parasite | —"— | Kurgan | 55.57 | 64.75 |
| A_HP_022 | 34 | 68 | 0.236 | Host-Parasite | —"— | Kustanai | 53.20 | 63.63 |
| A_HP_025 | 58 | 107 | 0.149 | Host-Parasite | —"— | Mongolia-North Western Khangay | 47.57 | 99.16 |
| A_HP_026 | 33 | 142 | 0.526 | Host-Parasite | —"— | Moscow | 55.75 | 37.62 |
| A_HP_027 | 47 | 108 | 0.212 | Host-Parasite | —"— | Moyyunkum | 44.28 | 72.95 |
| A_HP_029 | 49 | 79 | 0.155 | Host-Parasite | —"— | Kyrgyz Republic | 42.87 | 74.60 |
| A_HP_031 | 56 | 174 | 0.225 | Host-Parasite | —"— | Novosibirsk | 55.02 | 82.93 |
| A_HP_033 | 47 | 198 | 0.36 | Host-Parasite | —"— | Poland | 52.23 | 21.02 |
| A_HP_037 | 38 | 90 | 0.252 | Host-Parasite | —"— | Slovakia | 48.72 | 19.49 |
| A_HP_042 | 53 | 84 | 0.125 | Host-Parasite | —"— | Tarbagatai | 47.73 | 82.04 |
| A_HP_043 | 38 | 73 | 0.28 | Host-Parasite | —"— | Terskey Alatau | 42.15 | 78.44 |
| A_HP_044 | 53 | 197 | 0.281 | Host-Parasite | —"— | Tomsk-Tumen | 54.56 | 75.59 |

| ID | Focal Layer Species | Interaction Partners | Connectance | Type of interactions | Reference | Locality of Study | Latitude | Longitude |
|---|---|---|---|---|---|---|---|---|
| A_HP_046 | 56 | 202 | 0.305 | Host-Parasite | —"— | Turkmenistan | 39.59 | 58.95 |
| A_HP_047 | 37 | 100 | 0.35 | Host-Parasite | —"— | Tyva | 51.85 | 95.77 |
| A_HP_050 | 62 | 226 | 0.239 | Host-Parasite | —"— | Volga-Kama | 55.33 | 49.72 |
| A_HP_051 | 39 | 107 | 0.317 | Host-Parasite | —"— | Western-Sayan | 52.19 | 93.64 |
| M_PA_003 | 39 | 45 | 0.125 | Plant-Ant | Fonseca & Ganade (1996). Journal of Animal Ecology 66:339-347 | Biological Dynamics of Forest Fragments Project | -2.40 | -59.72 |
| M_PA_004 | 89 | 284 | 0.144 | Plant-Ant | Oikos 106:344-358 | Australian Canopy Crane | -16.12 | 145.45 |
| M_PL_006 | 78 | 146 | 0.141 | Pollination | Dicks et al. 2002. J. Anim. Ecol. 71:32-43. | Hickling, Norfolk, UK | 52.76 | 1.58 |
| M_PL_007 | 52 | 85 | 0.148 | Pollination | —"— | Shelfanger, Norfolk, UK | 52.41 | 1.10 |
| M_PL_017 | 104 | 299 | 0.151 | Pollination | Memmott 1999. Ecology Letters 2:276-280. | Bristol, England | 51.57 | -2.59 |
| M_PL_019 | 125 | 264 | 0.078 | Pollination | Inouye & Pyke. 1988. Australian Journal of Ecology 13:191-210. | Snowy Mountains, Australia | -36.45 | 148.27 |
| M_PL_025 | 57 | 143 | 0.25 | Pollination | Motten 1986. Ecological Monographs 56:21-42. | North Carolina, USA | 36.08 | -79.00 |
| M_PL_033 | 47 | 141 | 0.319 | Pollination | Small 1976. Canadian Field Naturalist 90:22-28. | Ottawa, Canada | 45.40 | -75.50 |
| M_PL_040 | 72 | 114 | 0.091 | Pollination | Ingversen (2006). Msc Thesis (Univ of Aarhus, Aarhus, Denmark). | Windsor, The Cockpit Country, Jamaica | 18.35 | -77.65 |
| M_PL_041 | 74 | 145 | 0.109 | Pollination | —"— | Syndicate, Dominica | 15.52 | -61.47 |

| ID | Focal Layer Species | Interaction Partners | Connectance | Type of interactions | Reference | Locality of Study | Latitude | Longitude |
|---|---|---|---|---|---|---|---|---|
| M_PL_045 | 43 | 63 | 0.143 | Pollination | Lundgren & Olesen (2005). Arc Antarc Alp Res 37:514-520. | Uummannaq Island, Greenland | 71.00 | -52.00 |
| M_PL_051 | 104 | 164 | 0.13 | Pollination | Vazquez 2002. Ph.D. Dissertation. University of Tennessee, Knoxville. | Nahuel Huapi National Park, Argentina | -41.08 | -71.53 |
| M_PL_058 | 113 | 319 | 0.123 | Pollination | Bartomeus et al. 2008. Oecologia 155: 761-770. | Parc Natural del Cap de Creus | 42.30 | 3.24 |
| M_PL_060_01 | 50 | 95 | 0.221 | Pollination | Kaiser-Bunbury et al. 2010. Ecology Letters 13:442-452. | Black River Gorges National Park, Mauritius | -20.70 | 57.73 |
| M_PL_060_02 | 50 | 107 | 0.235 | Pollination | —"— | —"— | -20.70 | 57.73 |
| M_PL_060_03 | 58 | 130 | 0.222 | Pollination | —"— | —"— | -20.70 | 57.73 |
| M_PL_060_04 | 67 | 134 | 0.139 | Pollination | —"— | —"— | -20.70 | 57.73 |
| M_PL_060_05 | 87 | 144 | 0.081 | Pollination | —"— | —"— | -20.70 | 57.73 |
| M_PL_060_06 | 71 | 96 | 0.082 | Pollination | —"— | —"— | -20.70 | 57.73 |
| M_PL_060_07 | 68 | 108 | 0.095 | Pollination | —"— | —"— | -20.70 | 57.73 |
| M_PL_060_08 | 47 | 72 | 0.135 | Pollination | —"— | —"— | -20.70 | 57.73 |
| M_PL_060_09 | 58 | 78 | 0.108 | Pollination | —"— | —"— | -20.70 | 57.73 |
| M_PL_060_10 | 39 | 51 | 0.146 | Pollination | —"— | —"— | -20.70 | 57.73 |
| M_PL_060_11 | 34 | 43 | 0.154 | Pollination | —"— | —"— | -20.70 | 57.73 |
| M_PL_060_12 | 37 | 60 | 0.21 | Pollination | —"— | —"— | -20.70 | 57.73 |
| M_PL_060_13 | 38 | 48 | 0.221 | Pollination | —"— | —"— | -20.70 | 57.73 |
| M_PL_060_14 | 48 | 99 | 0.243 | Pollination | —"— | —"— | -20.70 | 57.73 |
| M_PL_060_15 | 51 | 131 | 0.253 | Pollination | —"— | —"— | -20.70 | 57.73 |
| M_PL_060_16 | 56 | 114 | 0.172 | Pollination | —"— | —"— | -20.70 | 57.73 |
| M_PL_060_17 | 52 | 100 | 0.168 | Pollination | —"— | —"— | -20.70 | 57.73 |
| M_PL_060_18 | 48 | 75 | 0.134 | Pollination | —"— | —"— | -20.70 | 57.73 |

| ID | Focal Layer Species | Interaction Partners | Connectance | Type of interactions | Reference | Locality of Study | Latitude | Longitude |
|---|---|---|---|---|---|---|---|---|
| M_PL_060_19 | 31 | 41 | 0.175 | Pollination | —"— | —"— | -20.70 | 57.73 |
| M_PL_060_22 | 44 | 64 | 0.159 | Pollination | —"— | —"— | -20.70 | 57.73 |
| M_PL_060_23 | 39 | 57 | 0.176 | Pollination | —"— | —"— | -20.70 | 57.73 |
| M_PL_060_24 | 38 | 46 | 0.137 | Pollination | —"— | —"— | -20.70 | 57.73 |
| M_PL_061_05 | 34 | 51 | 0.193 | Pollination | Kaiser-Bunbury et al. 2014. Ecology, 95: 3314-3324. | Morne Seychellois National Park | -4.67 | 55.43 |
| M_PL_061_06 | 35 | 58 | 0.22 | Pollination | —"— | —"— | -4.67 | 55.43 |
| M_PL_061_07 | 32 | 62 | 0.3 | Pollination | —"— | —"— | -4.67 | 55.43 |
| M_PL_061_19 | 32 | 44 | 0.19 | Pollination | —"— | —"— | -4.67 | 55.43 |
| M_PL_061_23 | 36 | 45 | 0.173 | Pollination | —"— | —"— | -4.67 | 55.43 |
| M_PL_061_38 | 33 | 49 | 0.213 | Pollination | —"— | —"— | -4.67 | 55.43 |
| M_PL_061_40 | 35 | 58 | 0.248 | Pollination | —"— | —"— | -4.67 | 55.43 |
| M_PL_061_45 | 34 | 50 | 0.198 | Pollination | —"— | —"— | -4.67 | 55.43 |
| M_PL_061_46 | 34 | 45 | 0.165 | Pollination | —"— | —"— | -4.67 | 55.43 |
| M_PL_061_47 | 35 | 57 | 0.244 | Pollination | —"— | —"— | -4.67 | 55.43 |
| M_PL_063 | 64 | 123 | 0.248 | Pollination | Vizentin-Bugoni et al. (2016). Journal of Animal Ecology 85: 262-272. | Santa Virginia Field Station, Serra do Mar State Park | -23.34 | -45.12 |
| M_PL_066 | 36 | 73 | 0.471 | Pollination | Canela, M.B.F. (2006) Ph.D thesis. Universidade Estadual de Campinas, Brazil. | Serra da Mantiqueira, PNI, SE Brazil | -22.50 | -44.83 |
| M_PL_067 | 36 | 59 | 0.263 | Pollination | Las Casas et al. (2012) Brazilian Journal of Biology, 72, 51-58. | Serra do Para, Brazil | -7.87 | -36.40 |

| ID | Focal Layer Species | Interaction Partners | Connectance | Type of interactions | Reference | Locality of Study | Latitude | Longitude |
|---|---|---|---|---|---|---|---|---|
| M_PL_068 | 40 | 83 | 0.297 | Pollination | Gutierrez Zamora & Rojas Nossa (2001) BSc. Thesis. Universidad Nacional de Colombia, Colombia. | Santuario de Flora y Fauna Galeras | 1.25 | -77.43 |
| M_PL_071 | 52 | 89 | 0.253 | Pollination | Rosero (2003) Ph.D. Thesis. Universidade Estadual de Campinas, Brazil | Rainforest, Colombia | 0.04 | -72.27 |
| M_SD_002 | 40 | 119 | 0.427 | Seed Dispersal | Beehler 1983. Auk, 100: 1-12. | Mount Missim, Morobe Prov., New Guinea | -7.27 | 146.70 |
| M_SD_003 | 41 | 68 | 0.17 | Seed Dispersal | Carlo et al. (2003) Oecologia 134: 119-131 | Caguana, Puerto Rico | 18.30 | -66.78 |
| M_SD_004 | 54 | 95 | 0.14 | Seed Dispersal | —"— | Cialitos, Puerto Rico | 18.26 | -66.54 |
| M_SD_005 | 38 | 49 | 0.151 | Seed Dispersal | —"— | Cordillera, Puerto Rico | 18.17 | -66.59 |
| M_SD_006 | 36 | 51 | 0.162 | Seed Dispersal | —"— | Fronton, Puerto Rico | 18.31 | -66.56 |
| M_SD_010 | 64 | 234 | 0.334 | Seed Dispersal | Snow & Snow 1971. Auk, 88: 291-322. | Tropical rainforest. Trinidad. | 10.72 | -61.30 |
| M_SD_012 | 64 | 146 | 0.144 | Seed Dispersal | Galetti & Pizo 1996, Revista Brasileira de Ornitologia, 4: 71-79. | Santa Genebra Reserve T2. SE Brazil | -22.82 | -47.10 |
| M_SD_020 | 58 | 150 | 0.182 | Seed Dispersal | P. Jordano, unpubl. | Nava Correhuelas. S. Cazorla, SE Spain. | 37.93 | -2.87 |

| ID | Focal Layer Species | Interaction Partners | Connectance | Type of interactions | Reference | Locality of Study | Latitude | Longitude |
|---|---|---|---|---|---|---|---|---|
| M_SD_031 | 49 | 76 | 0.232 | Seed Dispersal | Ruben et al. (2013) Biological Invasions. 15, 5, 1143-1154. | Serra da Tronqueira | 37.80 | -25.19 |
| M_SD_034 | 121 | 419 | 0.139 | Seed Dispersal | Schleuning et al. (2011), Ecology, 92(1): 26-36. | Kakamega Forest, Kenya | 0.30 | 34.79 |
| Cambefort_Walter_1991 | 5 | 67 | 0.322 | Mammal-DungBeetle | Cambefort & Walter (1991). Dung beetles in tropical forests in Africa. In: Dung beetle ecology. [eds.: Hanski, I., Cambefort, Y.]. Princeton University Press | Makokou, Gabun | 0°33'44"N | 12°50'02.8"E |
| Slade_and_Raine_2016_EPT | 8 | 15 | 0.367 | Mammal-DungBeetle | E. Slade & E. Raine 2015/16, unpublished data | Serra do Mar, Paraná state, Brazil | 25°27'11.0"S | 48°52'57.0"W |
| Slade_and_Raine_2016_SPT | 8 | 19 | 0.401 | Mammal-DungBeetle | —"— | Serra do Mar, Paraná state, Brazil | 25°27'11.0"S | 48°52'57.0"W |
| Slade_and_Raine_2016_TPT | 8 | 18 | 0.368 | Mammal-DungBeetle | —"— | Serra do Mar, Paraná state, Brazil | 25°27'11.0"S | 48°52'57.0"W |
| Slade_and_Yuen_2016_Borneo_1 | 9 | 30 | 0.478 | Mammal-DungBeetle | Eleanor Slade & Li Yuen 2016/17, unpublished data | Luasong, Tawau, Sabah, Malaysia | 5°06'N | 117°30'E |
| Slade_and_Yuen_2016_Borneo_2 | 9 | 27 | 0.362 | Mammal-DungBeetle | —"— | Luasong, Tawau, Sabah, Malaysia | 5°06'N | 117°30'E |
| Slade_and_Yuen_Borneo_2016_3 | 9 | 35 | 0.565 | Mammal-DungBeetle | —"— | —"— | 5°06'N | 117°30'E |
| Slade_and_Yuen_Borneo_2016_4 | 8 | 33 | 0.527 | Mammal-DungBeetle | —"— | —"— | 5°06'N | 117°30'E |

| ID | Focal Layer Species | Interaction Partners | Connectance | Type of interactions | Reference | Locality of Study | Latitude | Longitude |
|---|---|---|---|---|---|---|---|---|
| Slade_and_Yuen _Borneo_2016_5 | 9 | 31 | 0.487 | Mammal-DungBeetle | —"— | —"— | 5°06'N | 117°30'E |
| Slade_and_Yuen _Borneo_2016_6 | 8 | 32 | 0.406 | Mammal-DungBeetle | —"— | —"— | 5°06'N | 117°30'E |
| Frank_2014_1 | 6 | 12 | 0.569 | Mammal-DungBeetle | Frank et al. (2017b) Agr. Ecosyst. Environ., 243:114-122. | Schorfheide, Germany (Exploratories) | 52°58'40.8"N | 13°44'50.6"E |
| Frank_2014_3 | 6 | 14 | 0.56 | Mammal-DungBeetle | —"— | Schorfheide, Germany (Exploratories) | 52°58'40.8"N | 13°44'50.6"E |
| Frank_2014_5 | 6 | 14 | 0.536 | Mammal-DungBeetle | —"— | Schwäbische Alb, Germany (Exploratories) | 48°25'32.0"N | 9°24'06.6"E |
| Frank_2014_7 | 6 | 17 | 0.461 | Mammal-DungBeetle | —"— | —"— | 48°25'32.0"N | 9°24'06.6"E |
| Frank_2015_13 | 6 | 14 | 0.571 | Mammal-DungBeetle | —"— | —"— | 48°25'32.0"N | 9°24'06.6"E |
| Frank_2015_19 | 12 | 13 | 0.346 | Mammal-DungBeetle | —"— | Hainich, Germany (Exploratories) | 51°10'00.3"N | 10°25'16.2"E |
| Hewavithana_2016 | 5 | 22 | 0.518 | Mammal-DungBeetle | Hewavithana et al. (2016). Int. J. Trop. Insect Sc., 36, 97-105. | Wasgomuwa National Park, Sri Lanka | 7°43'0"N | 80°56'0"E |
| Martin_Piera_Lobo_1996 | 8 | 35 | 0.457 | Mammal-DungBeetle | Martín-Piera & Lobo (1996) Miscell. Zoolog., 19, 13-31. | Coto de Doñana, National Park, Spain | 37°07'00.0"N | 6°27'00.0"W |
| Paetel_2002_South_Afric _Kruger-Park_natural_droppings | 6 | 85 | 0.573 | Mammal-DungBeetle | Paetel (2002). PhD Thesis, Humboldt-Universität zu Berlin, Germany. | Kruger park, South Africa | 24°59'00.0"S | 31°36'00.0"E |

| ID | Focal Layer Species | Interaction Partners | Connectance | Type of interactions | Reference | Locality of Study | Latitude | Longitude |
|---|---|---|---|---|---|---|---|---|
| Paetel_2002_South_Afric_Kruger-Park_pitfalls | 5 | 67 | 0.606 | Mammal-DungBeetle | Paetel (2002). PhD Thesis, Humboldt-Universität zu Berlin, Germany. | Kruger park, South Africa | 24°59'00.0"S | 31°36'00.0"E |
| Walter_1978_Congo | 8 | 96 | 0.613 | Mammal-DungBeetle | Walter (1978). Postdoctoral Thesis, Université des Sciences et Techniques du Languedoc, Montpellier, France. | Kinshasa, Congo | 5°06'31.0"S | 15°15'58.7"E |
| Whipple_Hoback_2012 | 11 | 15 | 0.842 | Mammal-DungBeetle | Whipple & Hoback (2012). Environ. Entomol., 41, 238-244. | Nebraska, USA | 41°28'08.4"N | 103°20'25.0"W |
| Wurmitzer_2017_1_corientes_A | 8 | 16 | 0.367 | Mammal-DungBeetle | Wurmitzer et al. (2017). Chemoecology, 27, 75-84.(Argentina) and unpublished data | Nacunan, Argentina | 34°02'42"S | 67°54'32"W |
| Wurmitzer_2017_2_corientes_B | 9 | 15 | 0.385 | Mammal-DungBeetle | —"— | Corrientes, Argentina | 28°01'40"S | 58°04'02"W |
| Wurmitzer_2017_4_nacunan_jan | 9 | 11 | 0.404 | Mammal-DungBeetle | —"— | Corrientes, Argentina | 28°01'40"S | 58°04'02"W |
| matrix_Abcoffeetotal | 9 | 5 | 0.289 | Host-Parasitoid | Tylianakis et al. (2007) Nature, 445:202-5 | | | |
| matrix_Coffeetotal | 12 | 7 | 0.262 | Host-Parasitoid | —"— | | | |
| matrix_Foresttotal | 10 | 5 | 0.3 | Host-Parasitoid | —"— | | | |
| matrix_Pasturetotal | 12 | 7 | 0.345 | Host-Parasitoid | —"— | | | |

| ID | Focal Species | Layer | Interaction Partners | Connectance | Type of interactions | Reference | Locality of Study | Latitude | Longitude |
|---|---|---|---|---|---|---|---|---|---|
| matrix_Ricetotal | 12 | | 7 | 0.31 | Host-Parasitoid | —"— | | | |
